# Inter-Subject EEG Correlation Reflects Time-Varying Engagement with Natural Music

**DOI:** 10.1101/2021.04.14.439913

**Authors:** Blair Kaneshiro, Duc T. Nguyen, Anthony M. Norcia, Jacek P. Dmochowski, Jonathan Berger

## Abstract

Musical engagement can be conceptualized through various activities, modes of listening, and listener states—among these a state of focused engagement. Recent research has reported that this state can be indexed by the inter-subject correlation (ISC) of EEG responses to a shared naturalistic stimulus. While statistically significant ISC has been reported during music listening, these reports have considered only correlations computed across entire excerpts and do not provide insights into time-varying engagement. Here we present the first EEG-ISC investigation of time-varying engagement *within* a musical work. From a sample of 23 adult musicians who listened to a cello concerto movement, we find varying levels of ISC throughout the excerpt. In particular, significant ISC is observed during periods of musical tension that build to climactic highpoints, but not at the highpoints themselves. In addition, we find that a control stimulus retaining envelope characteristics of the intact music, but little other temporal structure, also elicits significant neural correlation, though to a lesser extent than the original excerpt. In all, our findings shed light on temporal dynamics of listener engagement during music listening, establish connections between salient musical events and EEG ISC, and clarify specific listener states that are indexed by this measure.

## 1 Introduction

Musical engagement can implicate diverse aspects of the musical experience, from creation to interpretation to reception (Kaneshiro, 2016). Engaging with music is typically a hedonic experience able to evoke affective responses ranging from core affects to implicit empathy (Huron and Vuoskoski, 2020). Most studies of engagement focus on the listener, who can engage with music in a variety of ways including, among others, denotative, connotative, reflexive, and associative modes of listening (Huron, 2002). A broadly encompassing definition of listener engagement with music is suggested by Schubert et al. (2013) as being ‘compelled, drawn in, connected to what is happening, [and] interested in what will happen next’. In this paper we seek to identify this temporal process of attraction, attention, and anticipation by studying time-varying engagement with natural music across multiple listeners.

A number of studies have considered self-reports of musical engagement, including retrospective ratings delivered after listening to excerpts (Agres et al., 2017; Kaneshiro et al., 2020), questionnaires elucidating listeners’ general tendencies to enter states of engagement or absorption with music (Sandstrom and Russo, 2013), and continuous ratings delivered as music plays (Gregory, 1989; Broughton et al., 2019; Olsen et al., 2014). However, these approaches have known limitations. Retrospective reports and general listener profiles will not provide insights into specific stimulus events; and self-reports are subject to bias (Rosenman et al., 2011) and limited by a listener’s ability to self-assess (Madsen et al., 1993), and thus may be challenging or distracting to deliver—particularly continuously during music listening. On the other hand, a time-varying measure of engagement computed from responses delivered without effort by listeners—such as neural responses—could unobtrusively, and perhaps more objectively, index engagement with music as it is heard.

Electroencephalography (EEG) is a popular modality for studying neural processing of music, as its temporal resolution is sufficiently high to investigate specific musical events. Unfortunately, popular EEG paradigms involving repeated presentations of short, often tightly controlled stimuli tend to be incompatible with investigations of engagement involving complete naturalistic works in single stimulus exposures (Kaneshiro, 2016), and until recently, few studies (e.g., Leslie et al. (2014)) have used EEG to index engagement in realistic musical settings. However, inter-subject correlation (ISC) paradigms—in which audience members’ neural responses to a shared stimulus are correlated with one another—facilitate the study of engagement with single exposures to ecologically valid stimuli. The ISC approach was originally introduced as means of analyzing fMRI responses to naturalistic stimuli (Hasson et al., 2004); metholodogies for EEG were later introduced by Dmochowski et al. (2012). Since then, EEG studies using narrative stimuli—such as excerpts of films or speeches— have shown that ISC indexes narrative cohesion (Dmochowski et al., 2012; Ki et al., 2016), large-scale population preferences (Dmochowski et al., 2014), attentional state (Ki et al., 2016; Cohen et al., 2018), video viewership (Cohen et al., 2017), memory retention (Cohen and Parra, 2016), and even individual learning outcomes (Cohen et al., 2018). Given these findings, EEG ISC is sometimes interpreted as a measure of audience *engagement*, described by Dmochowski et al. (2012) as ‘emotionally laden attention’.

Processing of music has been investigated using ISC as well. fMRI studies have related the measure to neural tracking of time-varying stimulus features (Alluri et al., 2012), temporal cohesion of music (Abrams et al., 2013; Farbood et al., 2015), emotional responses to music (Trost et al., 2015), and effects of training (Fasano et al., 2020). Recent EEG studies, with a focus on engagement, have found ISC to be modulated by musical training, repeated exposures, and familiarity of musical genres (Madsen et al., 2019); and have used ISC and related measures to assess temporal stimulus manipulations, beat processing, and repetition effects during music listening (Kaneshiro et al., 2020).

One advantage of the high temporal resolution of EEG is that it permits ISC to be computed over short time windows, producing a time-varying measure of engagement on the scale of seconds. Dmochowski et al. (2012) applied this approach in their first EEG-ISC study of film viewing, finding that ISC not only varied over the course of a film excerpt, but also peaked during periods of high narrative tension and suspense. Cohen et al. (2017) later found that temporally resolved EEG ISC predicted real-world engagement with videos. As musical tension is thought to share the same underlying states as narrative tension in film excerpts— of dissonance and uncertainty seeking more stable ground (Lehne and Koelsch, 2015)—it is plausible that time-resolved ISC could highlight specific moments of high engagement during music listening as well. However, EEG-ISC studies of music listening to date have considered measures of neural correlation across entire excerpts only, on the order of minutes (Madsen et al., 2019; Kaneshiro et al., 2020). While these aggregate measures have yielded useful insights into music listening, they obscure temporal dynamics of musical engagement, which likely varies over the course of an excerpt. To the best of our knowledge, no investigation of time-varying musical engagement has yet been attempted using EEG ISC.

Temporally manipulated natural music has been used in neuroscience research for nearly 20 years (Levitin and Menon, 2003; Menon and Levitin, 2005). In the realm of ISC, both fMRI (Abrams et al., 2013) and EEG (Kaneshiro et al., 2020) studies have used phase-randomized (or *phase-scrambled*) music; this process preserves aggregate power spectra but disrupts the temporal structure of the audio. In both of these studies, phase-scrambled excerpts elicited lower neural correlation, and implicated different brain areas, than intact natural music. Thus, auditory stimulation alone is not thought to explain the neural correlation observed during music listening. However, as noted by both studies, phase scrambling removes numerous other structural elements of music, including but not limited to beat, phrase structure, instrumentation, and characteristic variations in loudness. Moreover, Kaneshiro et al. (2020) found that stimuli subjected to other, less extreme forms of temporal manipulation—i.e., stimulus reversal and shuffling at the measure (four-beat) level—elicited responses with even higher ISC than intact music. Therefore, there remain numerous stimulus attributes whose contributions to neural correlation have yet to be understood.

In the present study, we build off recent research and use EEG ISC to study time-varying engagement, for the first time with natural music. As stimuli we used a real-world musical work whose buildups to climatic events we consider conducive to Schubert et al. (2013)’s definition of engagement, in conjunction with a control stimulus providing incrementally more temporal structure than phase-scrambled stimuli used previously (Abrams et al., 2013; Kaneshiro et al., 2020). We took an exploratory approach based on past investigations of temporally resolved EEG ISC during film viewing (Dmochowski et al., 2012) and conducted analyses over three maximally correlated EEG components. We hypothesized first that ISC would vary over the course of the original excerpt and peak at preselected musically salient events such as buildups to musical ‘highpoints’ (Agawu, 1984), which we consider analogous to periods of tension and suspense in more explicitly narrative stimuli such as films (Dmochowski et al., 2012; Lehne and Koelsch, 2015). Second, based on the importance of the stimulus amplitude envelope in speech processing (Van Tasell et al., 1987; Drullman et al., 1994; Shannon et al., 1995; van der Horst et al., 1999; Aiken and Picton, 2008), we hypothesized that our control stimulus—which applies the envelope of the intact music onto a phase-scrambled version—would elicit responses with higher neural correlation than phase scrambling alone. However, we also predicted that fluctuations in audio amplitude envelope would play a role in, but not fully explain, neural correlation during music listening, and thus expected that intact music would elicit higher ISC.

## 2 Methods

### 2.1 Ethics Approval Statement

This research was approved by the Institutional Review Board at Stanford University (protocol number IRB-28863). All participants delivered written informed consent prior to participating in the experiment

### 2.2 Data Availability Statement

The data that support the findings of this study (i.e., raw, continuous EEG recordings; preprocessed, aggregated EEG trials; and behavioral ratings with anonymized participant identifiers) are openly available from the Stanford Digital Repository as part of the Natural-istic Music EEG Dataset—Elgar (NMED-E) at https://purl.stanford.edu/pp371jh5722 (Kaneshiro et al., 2021).

### 2.3 Stimuli

#### 2.3.1 Stimulus Selection and Characterization

We used two stimuli in this study: The complete opening movement of Edward Elgar’s Cello Concerto in E minor, Op. 85, performed by Jacqueline du Pré and the London Symphony Orchestra, conducted by Sir John Barbirolli; as well as a temporally manipulated version of the performance as described below. Composed in 1919, Elgar’s concerto is a dramatic work with a wide dynamic and expressive range, and highly contrasting textures. The work comprises two waves of dramatic gestures that sweep to culminating ‘highpoints’ (Agawu, 1984), which then lead to structural segmentation boundaries. du Pré’s influential 1965 recording is known for reviving the work and bringing it into the standard orchestral repertoire (Solomon, 2009). The concerto, and in particular the recording we chose, is widely noted as being highly expressive and arousing.^1^

The movement of the concerto is in ternary form with the framing sections in E minor, and the middle section primarily in E major. The sections are linked through two similar motivic elements, which both share a long-short-long rhythmic pattern encompassing neighboring melodic motion. The unifying motivic variants pervade the movement varying in tempi, underlying harmony, melody, orchestration, and with highly contrasting levels of intensity. The framing A and A’ sections each include a climactic event preceded by the main theme played by the soloist, followed by textural and dynamic increases that culminate in a melodic ascent to an E-minor cadence.

To guide our data analysis, we identified in advance a set of salient musical events (shown as dashed vertical lines in Figure 1A–B), which we conjectured would elicit significantly correlated ISC responses. First, we labeled as E1 and E1’ the two introductions of the main theme played by the soloist, which signify the start of each buildup to climactic highpoints in the movement. The theme is then taken up by the strings at E2 and E2’, and in E3 is played by the soloist, now definitively in the tonic key of E minor (E3 is not reprised in the second buildup). Finally, an ascending melodic line played by the soloist reaches a climactic E-minor cadence at E4 and E4’, which are the structural highpoints. We predicted that ISC would be significant between E1 and E4—and later between E1’ and E4’—because these are regions of rising tension and suspense which ultimately culminate in the cadential highpoints (cf. Schubert et al. (2013)’s definition of engagement and ISC peaks reported by Dmochowski et al. (2012)). Finally, we labeled as E5 the start of the B section of the movement. Here we anticipated that the cessura at the point of arrival and subsequent introduction of new musical themes would elicit high ISC.

**Figure 1:**
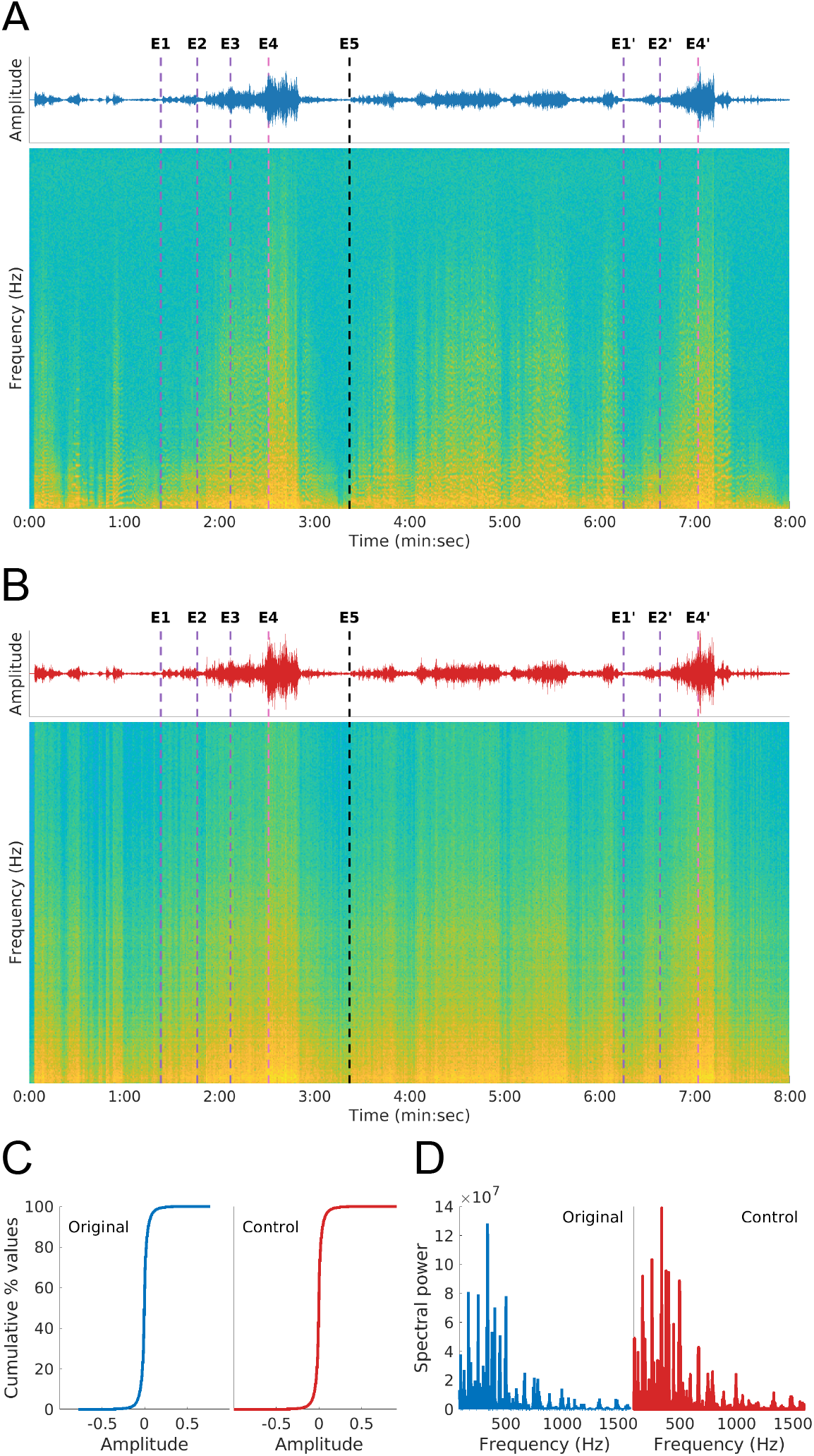
Experimental stimuli. (**A**) Audio waveform and spectrogram of the natural music (Original) stimulus, an influential recording of the first movement of Elgar’s Cello Concerto (1919), performed in 1965 by Jacqueline du Pré and the London Symphony Orchestra, conducted by Sir John Barbirolli. Dashed vertical lines denote preselected stimulus events of interest: Instances of the main theme played by the solo cello (E1, E1’), strings (E2, E2’), and again by the soloist (E3), which lead to structural highpoints (E4, E4’); and the start of second section in the movement (E5). (**B**) Audio waveform and spectrogram of the Control stimulus: A phase-scrambled version of the Original stimulus which was subsequently scaled by the amplitude envelope of the intact music. Dashed lines are as described in (A). (**C**) Cumulative distributions of waveform values are similar across stimuli. (**D**) Stimuli share similar aggregate power spectra.

#### 2.3.2 Stimulus Preparation

Both stimuli were 480 seconds (8 minutes) in length. We purchased the EMI Records, Ltd. digitally remastered version of the recording (Elgar et al., 1999) in digital format from iTunes. Using Audacity^2^ recording and editing software, we remixed the .m4a stereo recording to mono, and exported the result to .wav format. Subsequent audio processing was performed using Matlab^3^ software. First, a linear fade-in and fade-out were applied to the first and last 1 second of the recording, respectively; the resulting audio served as the *Original* stimulus.

The audio was processed further to create the *Control* stimulus. First, we performed the phase-scrambling procedure described in Kaneshiro et al. (2020), converting the audio waveform to the frequency domain, randomizing phase values at each frequency bin while preserving conjugate symmetry, and converting the result back to the time domain. Following this, we used the publicly available MIRtoolbox (Lartillot and Toiviainen, 2007) to compute the audio envelope of the Original stimulus, and subsequently scaled the phase-scrambled audio by this envelope. The result is reminiscent of envelope-shaped noise stimuli that have been used in past speech studies (Horii et al., 1971; Van Tasell et al., 1987; Shannon et al., 1995). Finally, we scaled the entire Control waveform so that its global RMS equaled that of the Original.

Both stimuli are visualized in Figure 1. As shown in Panels A and B, Original (blue) and Control (red) waveforms are visually similar but not identical. We also visualized the stimuli as spectrograms,^4^ which highlight spectral harmonics in the Original but not the Control stimulus (e.g., around 1:00). While phase scrambling alone preserves the exact power spectrum of the original audio, the additional envelope-scaling procedure introduces a similar but not identical power spectrum. Panels C and D show that the two stimuli remain similar on the basis of cumulative distributions of waveform values (computed from histograms with 500 bins each) and aggregate power spectra, respectively.

### 2.4 Experimental Paradigm

The data analyzed here are part of a larger study on multimodal listener responses to natural music. The larger study comprised a demographic and music experience questionnaire, the Absorption in Music Scale (AIMS) questionnaire (Sandstrom and Russo, 2013), and two experimental blocks of music listening. Each participant’s data were collected in a single experimental session: After the participant delivered informed consent, they filled out the questionnaires and then completed the music-listening blocks. In the first block, after 194 completing a short training session to become familiarized with the experimental and task paradigms, the participant heard each stimulus once while EEG, electrocardiogram (ECG), and chest and abdomen respiratory plethysmography were recorded simultaneously. In order to avoid biasing responses, participants were not prompted with a definition of engagement in this block. Each stimulus was preceded by a 60-second ‘baseline’ period to be used during analysis of physiological responses. During this interval, pink noise was played at a low volume while the participant sat still with eyes open and performed no task. Each baseline period was followed by a stimulus, to which the participant listened attentively with eyes open while avoiding movement and viewing a fixation image shown on a monitor 57 cm in front of them. After each stimulus, participants delivered key-press ratings on a scale of 1 to 9 on the degree of pleasantness, arousal, level of interest, predictability, and familiarity of the stimulus just heard; these questions, adapted from those used in a previous EEG-ISC music study (Kaneshiro et al., 2020), are intended to probe aspects of music listening relating to both engagement and (for the larger scope of the study) arousal. After the intact excerpt only, participants also used the same response scale to report how often they listened to the genre of classical music. The EEG electrode net and physiological sensors were removed after the first block. In the second block, the participant used a mouse-operated onscreen slider to report their level of engagement over time with each stimulus as it played; here they were guided by Schubert et al. (2013)’s definition of engagement. In each block, stimuli were presented in a pseudo-random order ensuring an equal distribution of orderings across the original 24 participants. Our present analysis considers demographic information as well as the EEG data and behavioral ratings collected in the first experimental block; thus, further descriptions of data acquisition, preprocessing, and analysis will focus only on these responses.

### 2.5 Participants

We recruited right-handed participants who were 18–35 years of age, had normal hearing, had no cognitive or decisional impairments, and were fluent in English. As past research has shown that formal musical training is associated with enhanced cortical responses to music (Pantev et al., 1998), we sought participants who had at least five continuous or non-continuous years of formal training in classical music; this could include private lessons in instrumental or vocal performance, composition lessons, and AP- or college-level music theory courses. We recruited participants who enjoyed listening to classical music at least occasionally, but sought participants with no cello experience in order to avoid involuntary motor activations that can occur when hearing music played by one’s own instrument (Haueisen and Knösche, 2001). All participants confirmed their eligibility prior to participating.

From 24 participants who completed the experiment, one participant’s data were excluded during preprocessing—prior to spatial filtering and ISC analyses—due to gross noise artifacts in the EEG recording (see § 2.6). We thus obtained usable data from *N* = 23 participants (7 male) ranging in age from 19 to 33 years (*m* = 23.26 years, *s* = 4.73 years). From this sample, years of musical training ranged from 5 to 18 years (*m* = 10.70 years, *s* = 3.47 years), and classical-music listening was confirmed through both the verbal eligibility confirmation and a behavioral report delivered during the experiment (see Figure 4).

### 2.6 Data Acquisition and Preprocessing

Stimulus presentation and key-press response acquisition were programmed using Matlab’s Psychophysics Toolbox (Brainard, 1997). Other aspects of stimulus delivery and EEG data acquisition were as described in Kaneshiro et al. (2020): Mono sound files were played through two magnetically shielded Genelec 1030A speakers, which were located 120 cm from the participant in an acoustically and electrically shielded ETS-Lindgren booth. The second audio channel (not played to participants) contained intermittent square-wave pulses, which were sent directly to the EEG amplifier for precise time stamping of stimulus onsets. EEG was recorded from 128-channel nets using the Electrical Geodesics (EGI) GES 300 platform (Tucker, 1993) and Net Station software at a sampling rate of 1 kHz, referenced to the vertex. Responses to both stimuli, their preceding baseline intervals, delivery of ratings, and a short break after the first stimulus trial were acquired in a single recording. Electrode impedances were no higher than 60 kΩ at the start of each EEG recording (Ferree et al., 2001).

Continuous EEG recordings were exported to Matlab file format using Net Station software. Subsequent preprocessing and analysis were performed on a per-recording basis, using custom Matlab code as well as publicly available third-party code as described below. First, the continuous recording was highpass (0.3 Hz, 8th-order Butterworth), notch (59–61 Hz, 8th-order Butterworth), and lowpass (50 Hz, 8th-order Chebyshev) filtered using zero-phase filters, and then downsampled by a factor of 8 to a final sampling rate of 125 Hz. Following this, we extracted baseline and stimulus trial labels; event triggers of timing clicks sent from the audio; and behavioral ratings. Baseline and stimulus trials were then epoched using the precise timing triggers sent from the audio: Baseline trials (which were preprocessed but are not analyzed here) were 60 seconds (7501 time samples) in length, and stimulus trials were 480 seconds (60001 time samples) in length. We performed a median-based DC correction on each of the four epochs and concatenated them in time. EOG channels were computed from electrodes above and/or below the eyes (VEOG) and at the sides of the eyes (HEOG). We retained electrodes 1–124 for further analysis, excluding electrodes on the face; bad electrodes—identified during the experimental session based on impedances and/or by visual inspection of the raw data, or with at least 10% of voltage magnitudes exceeding 50 *μ*V across the recording after ICA artifact removal—were also removed at this point, reducing the number of rows in the data matrix. We used the EEGLAB Toolbox (Delorme and Makeig, 2004) implementation of extended Infomax ICA to identify and remove ocular and ECG artifacts from the data in a semi-automatic fashion (Bell and Sejnowski, 1995; Jung et al., 1998): Ocular artifacts were removed according to EOG correlation thresholds described in Kaneshiro et al. (2020), while ECG artifacts were removed by visual inspection of the first 30 components’ time-domain waveforms. Subsequent preprocessing steps are as described in Kaneshiro et al. (2020), including identification and exclusion of recording- and trial-wide bad electrodes; replacement of noisy transients with NaNs; replacement of missing rows from bad electrodes with rows of NaNs (ensuring the same number of rows in each data trial); addition of the vertex reference channel and conversion of the data matrix to average reference; replacement of NaNs with the average of data from neighboring electrodes; and a final, mean-based DC correction. During the preprocessing stage, one participant was excluded from further analysis due to gross noise artifacts throughout their recording.

Following preprocessing of all recordings, the trials for each stimulus were aggregated across participants. This produced an electrode-by-time-by-trial matrix for each stimulus, the size of which was 125 × 7501 × 23 for baseline stimuli and 125 × 60001 × 23 for Original and Control stimuli.

### 2.7 Analysis of EEG Data

We analyzed EEG responses to the Original and Control stimuli (excluding the baseline responses) on a per-stimulus basis, using a two-step process first introduced by Dmochowski et al. (2012). First, the EEG data matrices were factorized to compute three spatial components in which ISC was maximized. Following this, we computed ISC across entire excerpts 288 and in a time-resolved fashion.

#### 2.7.1 Reliable Components Analysis (RCA)

As preparation for the ISC calculations, we first spatially filtered the data—that is, we computed optimal spatial ‘components’ (linear combinations of electrodes) in which ISC of the projected EEG data would be maximized. For this step, we used Reliable Components Analysis (RCA), which was first introduced in the EEG-ISC study by Dmochowski et al. (2012). This procedure concentrates the most-correlated activity across trials—which might be spatially distributed across the electrode montage—into a small number of components. In the case of time-domain analyses as used here, RCA provides ISC-optimized temporal responses while reducing the spatial dimensionality of the data from 125 electrodes to three components. RCA is similar to PCA in that it involves an eigenvalue decomposition; therefore, the procedure returns multiple component weight vectors (eigenvectors)—which for RCA are sorted in descending order of ISC—and accompanying coefficients (eigenvalues). Following Dmochowski et al. (2012), we analyzed the first three Reliable Components (RC1–RC3).

Therefore, each trial of data—a time-by-electrode matrix (60001 × 125)—was multiplied by an electrode-by-component (125 × 3) weight matrix to produce a time-by-component matrix (60001 × 3). ISC was then computed using each component vector (60001 × 1).

We used a publicly available Matlab implementation (Dmochowski et al., 2015)^5^ to perform RCA using the procedure and parameters reported in Kaneshiro et al. (2020). RCA components were computed separately for each stimulus. As reported in Kaneshiro et al. (2020), we visualized individual components as scalp topographies using forward-model projections of the weight vectors (Haufe et al., 2014; Parra et al., 2005) and report the scalar coefficient of each component.

#### 2.7.2 Inter-Subject Correlation (ISC)

ISC was computed across participants on a per-RC, per-stimulus basis. Each ISC calculation involved *N* = 23 single trials of time-domain response data from a single RC for a single stimulus. Each participant’s ISC was computed as the mean correlation of their trial with all other trials. Following that, we computed mean ISC, and standard error of the mean, across participants. We computed ISC in two ways: First, over the entire duration of the stimulus, as was done in past music studies (Madsen et al., 2019; Kaneshiro et al., 2020), and second, in a temporally resolved fashion. For the latter case we followed the procedure of Dmochowski et al. (2012), computing ISC over a 5-second window that advanced in 1-second increments. This produced an ISC time course with a temporal resolution of 1 second, totalling 476 time windows for each 480-second stimulus.

### 2.8 Statistical Analyses

Statistical significance of the EEG results was computed in two ways. First, we assessed the standalone statistical significance of each RCA coefficient and ISC using permutation testing. As described in Kaneshiro et al. (2020), this involved 1,000 RCA and subsequent ISC calculations for each stimulus; in each calculation, every trial of data input to RCA was first phase scrambled, which disrupts across-trials temporal covariance while retaining aggregate power spectra and autocorrelation characteristics of individual trials (Prichard and Theiler, 1994). This produced a null distribution of 1,000 surrogate observations for each analysis, the 95th percentile of which served as the threshold for statistical significance. In reporting the significance of RCA coefficients, as well as ISC computed across entire stimulus durations, we corrected for multiple comparisons using False Discovery Rate (FDR; Benjamini and Yekutieli (2001)). For this we report adjusted *p*_*FDR*_ values, corrected across 3 comparisons (components) on a per-stimulus basis. We present the time-resolved ISC for each stimulus and RC in relation to the 95th percentile of its null distribution, which is also a time series. Here we do not perform any cluster correction, as temporal dependencies are accounted for through the preservation of autocorrelation in the 1,000 phase-scrambled records (Theiler et al., 1992; Prichard and Theiler, 1994; Lancaster et al., 2018).

The second form of statistical analysis involved across-condition comparisons of ISC and behavioral ratings. For ISC, we focused on RC1 results only, as this was the only component with statistically significant ISC for both stimulus conditions. For the behavioral ratings and RC1 ISC computed across entire stimulus durations, we performed paired, two-tailed Wilcoxon signed-rank tests across the distributions of *N* = 23 participants. As behavioral responses to multiple questions were given, we corrected for 5 comparisons and report *p*_*FDR*_ values. Next, we wished to know whether the temporally resolved RC1 ISC differed significantly overall according to stimulus. To test this, we performed a paired, two-tailed Wilcoxon signed-rank test across the two vectors of across-participant mean ISC values (*N* = 476 time points per vector). Finally, we correlated the group-mean RC1 ISC time series across stimulus conditions to determine whether their activations were similar, even if their overall level differed by condition.

## 3 Results

### 3.1 Original and Control Stimuli Elicit Correlated Components

We used inter-subject correlation (ISC) to analyze EEG responses to an intact excerpt of natural music and a temporally manipulated control. Prior to computing ISC, we spatially filtered the data, projecting EEG trials from electrode space to three components in which 356 ISC was maximized. The RC1–RC3 component topographies, along with their coefficients and ISC computed across entire durations of stimuli, are shown in Figure 2. As shown in Panel A of the figure, the first component (RC1) is broadly similar across stimulus conditions, with a fronto-central topography.^6^ Panel B shows the RCA coefficients (eigenvalues) for the first three RCs in line plots, with the respective 95th percentile of the permutation testing null distribution represented as the height of each shaded gray area. As suggested by these plots, the RC1 coefficient is statistically significant for the Original, but not the Control stimulus (permutation test, Original RC1 *p*_*FDR*_ < 0.001, Control RC1 *p*_*FDR*_ = 0.23, corrected for 3 comparisons per condition). The topographies for RC2 and RC3 differ by stimulus condition, though their coefficients are not statistically significant (all *p*_*FDR*_ ≥ 0.23).

**Figure 2:**
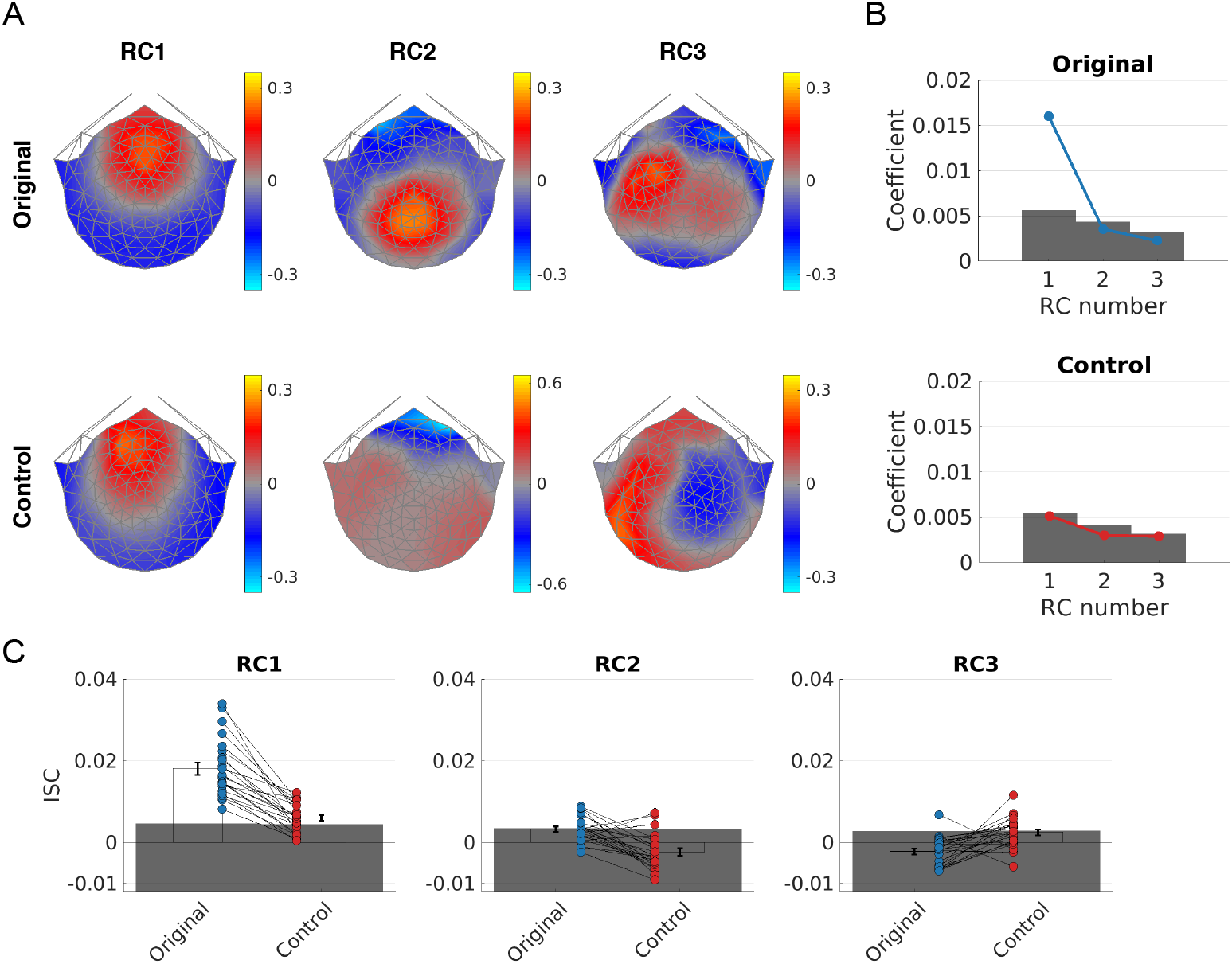
Aggregate measures of neural correlation. (**A**) Forward-model projected topographies of the three maximally correlated components (RC1–RC3), computed separately for each stimulus. (**B**) RC1–RC3 coefficients. Height of the shaded gray area denotes the 95th percentile of the null distribution for each component. Only the RC1 coefficient for the Original stimulus is statistically significant (permutation test, FDR corrected). (**C**) RC1–RC3 ISC computed across the entire duration of each stimulus. Error bars represent *±* SEM, and points represent individual participants. Height of the shaded gray area denotes the 95th percentile of the null distribution for each group-level ISC. RC1 ISC is statistically significant for both Original and Control stimuli (permutation test, FDR corrected), and differs significantly between stimuli (Wilcoxon signed-rank test) with higher ISC for the Original excerpt. RC2 and RC3 ISC were not significant for either stimulus (permutation test, FDR corrected), and thus were not compared across conditions.

### 3.2 Aggregate ISC Is Higher for Intact Music

We computed ISC of RC1–RC3 data across the entire duration of each stimulus; per-component, per-condition results are shown in Figure 2C. For RC1, mean ISC exceeds the 95th percentile of the null distribution (upper edge of shaded gray area) for both stimuli (permutation test, Original and Control *p*_*FDR*_ < 0.001, corrected for 3 comparisons per condition). A follow-up paired comparison between stimulus conditions indicates that RC1 ISC differs significantly according to stimulus condition (two-tailed Wilcoxon signed-rank test, *W* = 276, *p* < 0.001), with higher mean ISC for the Original stimulus. However—perhaps unsurprisingly given the RCA coefficients (Panel B)—all-time ISC is not significant in RC2 or RC3 for either stimulus (all *p*_*FDR*_ ≥ 0.12); therefore, they are not analyzed further.

### 3.3 Temporally Resolved ISC Implicates Salient Musical Events

For each stimulus and RC component, we next computed ISC over 5-second windows, with a temporal resolution (window shift size) of 1 second. Results are shown in Figure 3. As can be seen in Panel A, RC1 ISC of the Original stimulus often exceeds the permutation testing significance threshold (top of shaded gray area), with a total of 37.18% significant time windows (permutation test, uncorrected; see § 2.8). As shown in Panels B and C, however, time-varying ISC rarely reaches statistical significance for RC1 of the Control stimulus, or for RC2 or RC3 of either stimulus.

**Figure 3:**
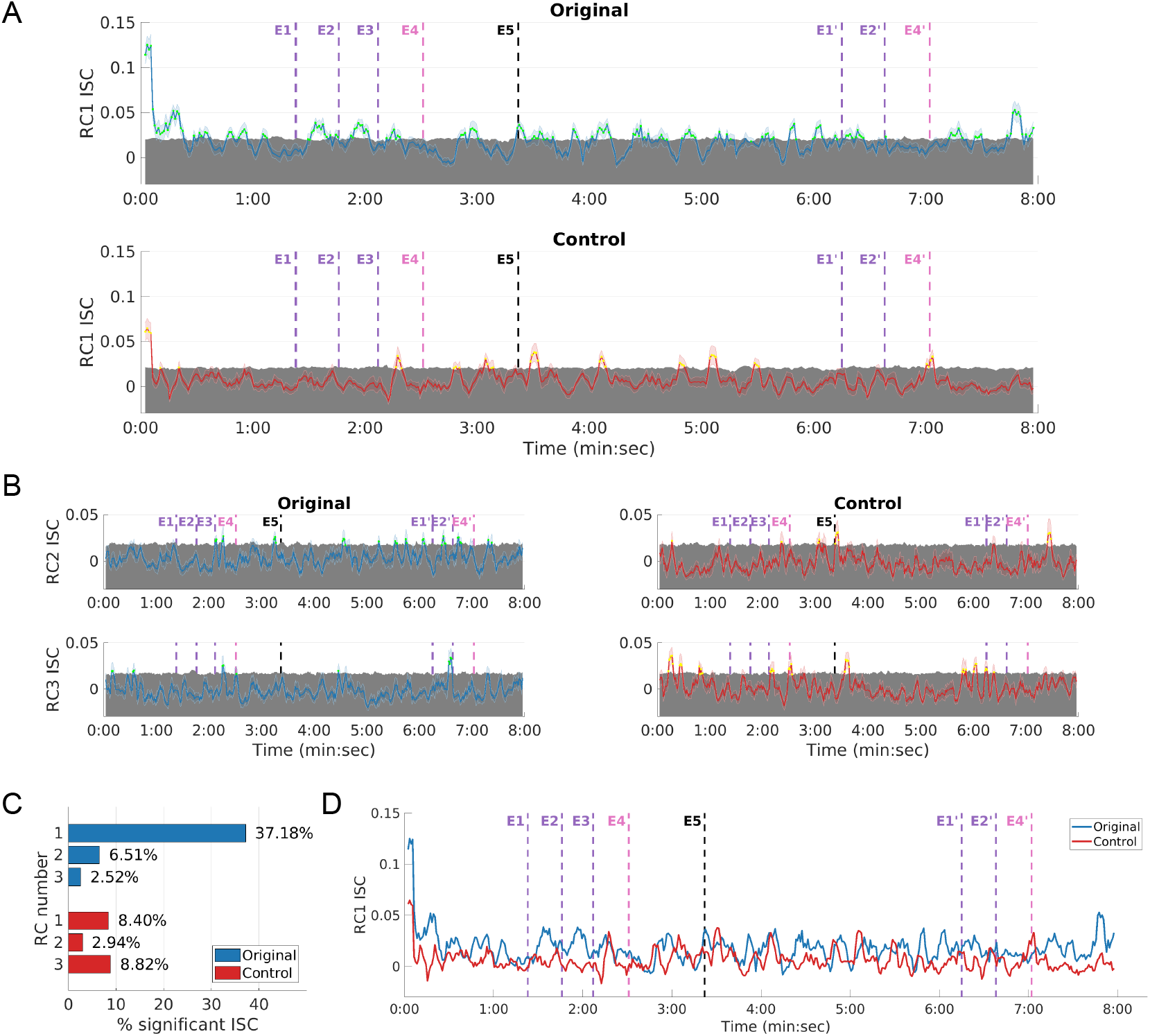
Time-varying ISC. ISC was computed in 5-second windows advancing in 1-second increments (colored lines) and plotted in relation to the 95th percentile of the null distribution (shaded gray area). Dashed vertical lines denote musically salient events (see Figure 1). Statistically significant ISC (permutation test, uncorrected) is denoted in green (Original stimulus) and orange (Control stimulus). (**A**) Temporally resolved ISC for RC1, with line plots representing mean ISC and error bars representing *±* SEM. (**B**) Temporally resolved ISC for RC2 and RC3, with line plots representing mean ISC and error bars representing *±* SEM. (**C**) Percentage of time windows containing statistically significant ISC for each stimulus and RC 1–3 (permutation test, uncorrected). (**D**) Overlay of mean RC1 ISC for each condition. The Original stimulus elicits higher time-resolved RC1 ISC than the Control (Wilcoxon signed-rank test), and time courses are not strongly correlated (*r* = 0.357, all time windows; *r* = 0.050, first four time windows excluded).

**Figure 4:**
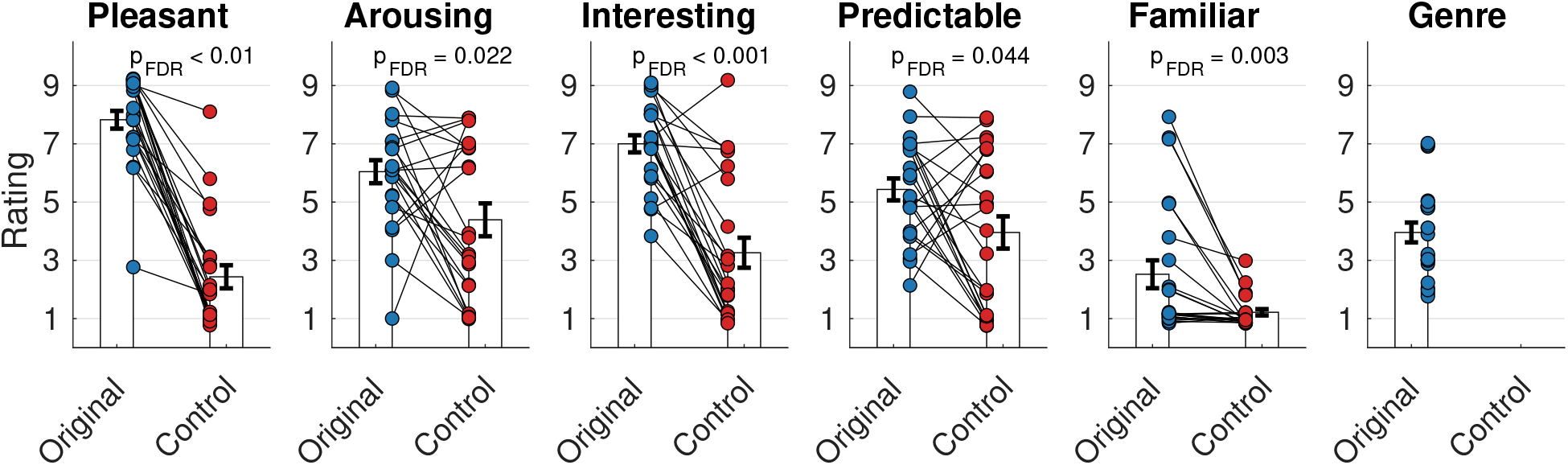
Behavioral ratings. Participants rated each stimulus on a 1–9 ordinal scale along dimensions of pleasantness, arousal, level of interest, predictability, and familiarity; and reported how often they listen to classical music. Bar heights represent mean values, error bars represent *±* SEM, and dots represent individual responses. Ordinal responses (y-values) are slightly jittered for visualization only. All between-condition comparisons were statistically significant (Wilcoxon signed-rank test, FDR corrected), with the Original stimulus receiving higher ratings for each dimension. All participants listened to classical music at least occasionally.

We observe significant ISC around some but not all preselected points in the performance: In the first leadup to the highpoint at E4, ISC is significant after each demarcated event (E1, E2, E3). ISC is also significant at the start of the second section (E5). However, ISC is lower overall in the shorter second leadup (E1’, E2’) to the highpoint at E4’, and is not significant at either highpoint arrival (E4, E4’). Temporally resolved ISC was significant at numerous additional points during the Original excerpt. Other notable peaks occur during the first 21 and last 15 ISC windows (Figure 3). The former peaks highlight the opening portion of the excerpt, which contains an extended dramatic solo passage that establishes the key of the movement. The latter peak consists of the solo cello, followed by the strings, reprising the sparse closing cadence of the movement’s first section (leadup to E5).

In a post-hoc analysis, we compared the percentage of significant ISC windows corresponding to the leadups to the first highpoint (E1 to E4) and the second highpoint (E1’ to E4’), since these are the regions of rising tension and suspense. 44.29% of the ISC windows were significant in the leadup to the first highpoint, while 21.28% were significant in the leadup to the second highpoint.

Figure 3D shows the overlaid RC1 time-resolved ISC for both stimulus conditions (as shown in Panel C, RC1 was the only component for which the percentage of significant ISC windows exceeded the *α* level of 0.05 for both stimuli). As suggested by the percentages of significant RC1 ISC reported for each condition, the distributions of temporally resolved ISC differed according to stimulus condition (two-tailed Wilcoxon signed-rank test, *W* = 99342, *p* < 0.001), with higher overall ISC for the Original condition. Finally, we note that while both stimuli elicited high ISC in the early time windows, the two RC1 ISC time series are not highly correlated (*r* = 0.3572 over all time windows; *r* = 0.0497 when the first four time windows are excluded).

### 3.4 Behavioral Ratings Are Higher for Intact Music

Participants reported how pleasant, arousing, interesting, predictable, and familiar they found each stimulus on an ordinal scale of 1 to 9. Ratings are shown in Figure 4; y-values have been slightly jittered for visualization purposes only. Responses differed significantly according to stimulus for all questions (two-tailed Wilcoxon signed-rank test, all *W ≥* 55, all *p*_*FDR*_ ≤ 0.044, corrected for 5 comparisons), with higher mean responses for the Original stimulus in all cases. Participants also reported how often they listened to the genre of the Original stimulus (i.e., classical music); all responded higher than 1 (‘Never’), confirming this aspect of their eligibility.

## 4 Discussion

EEG inter-subject correlation (ISC) has been found to index states of engagement with naturalistic stimuli such as film excerpts, speeches, and musical works. In this study we have for the first time analyzed temporally resolved EEG ISC in the context of music listening. Through this approach we have confirmed that the ISC measure of engagement varies over the course of a real-world musical work, as was previously shown for film viewing (Dmochowski et al., 2012; Poulsen et al., 2017). Moreover, in extending a phase-scrambling stimulus manipulation used in past music ISC studies (Abrams et al., 2013; Kaneshiro et al., 2020), we have shown that an audio amplitude envelope characteristic of music elicits correlated EEG responses, though to a lesser extent than intact music.

EEG responses to film excerpts contained ISC peaks during periods of high tension and suspense (Dmochowski et al. (2012, Fig. 1); replicated by Poulsen et al. (2017, Fig. 2) with mobile EEG). Based on these findings, we expected ISC to vary during music listening as well, and to peak during periods in which the music induced a state of *engagement* as defined by Schubert et al. (2013): Being ‘compelled, drawn in, connected to what is happening, [and] interested in what will happen next’. Indeed, across an audience of 23 adult musicians listening to the first movement of Elgar’s Cello Concerto, we observed temporal variations in EEG ISC with a number of significant peaks (Figure 3).

In designing the study, we identified events in the intact stimulus that we expected would correspond to high ISC based upon the dramatic intensity of the music (see § 2.3.1). Some but not all of these events corresponded to significant ISC. In the first climactic rise to the structural highpoint (E1–E4), ISC was significant during periods of escalating tension leading up to the highpoint itself; this is confirmatory with previous findings relating to film viewing (Dmochowski et al., 2012). However, at each highpoint’s arrival, ISC was not only nonsignificant but was so for extended periods of time relative to the rest of the excerpt. This lack of significant ISC at and following the very moments which are deemed climactic may correspond to a conspicuous absence of musicological, music-theoretic, and psychological studies focusing on the affective processes from a point of maximal tension until the actual resolution of tension. Most literature treats this transition as a binary switch in which tension is suddenly resolved. Music theorists have sought to nuance the process delineating patterns of fluctuating intensity leading to and following a climax in terms of intensity (Berry, 1978) or narrative (Childs, 1977; Agawu, 1984); the affective peaks and troughs of these processes are described in terms of intensification, climax, and abatement (Patty, 2009). There is, however, a considerable lack of consistency and a wide variety of issues, terminology, and analytical approaches, and thus this key aspect of musical engagement begs further study.

While we know of no EEG-ISC study focusing specifically on moments of apotheosis (i.e., the culminating moment of tension and suspense) during film viewing, we posit that our finding of low ISC at the resolution of suspense reflects a state distinct from the preceding, increasingly anticipatory state. We do not suggest that listener engagement ceases or decreases at the musical climax, but rather that the specific nature of engagement at this point is not evident or indexed by ISC. Recent work by Chabin et al. (2020) has used EEG oscillatory band power to identify neural correlates of chills and emotional pleasure during natural music listening, while Czepiel et al. (2020) linked ISC of physiological responses to live music to musical elements such as structural segmentation boundaries. Thus, future work could consider using EEG-ISC alongside complementary analysis approaches and response modalities to more fully characterize listener experiences of music.

RC1 ISC peaks were observed at the beginning of both the Original and Control stimuli (Figure 3A); these are likely partly attributable to low-level processing of the stimulus onset. However, ISC is markedly higher here for the Original stimulus compared to the Control, with peak correlations of 0.13 and 0.06, respectively. In addition, the Original but not the Control stimulus elicited a second sustained peak at around 19 seconds. Given the musical content of this section (see § 3.3), we interpret these peaks as reflecting not only low-level processing but also listeners being ‘drawn in’ to the music and ‘interested in what will happen next’, consistent with Schubert et al. (2013)’s definition of engagement.

The ternary structure of the Original stimulus enabled us to compare ISC across repeated periods of musical tension and suspense. The larger percentage of significant ISC windows in the leadup to the first highpoint compared to the second suggests that repetition effects—in which EEG ISC drops over repeated exposures (Dmochowski et al., 2012; Poulsen et al., 2017; Madsen et al., 2019; Kaneshiro et al., 2020)—may also occur locally within a work. Yet we see contradictory results as well, as repeated material appearing first before E5 and again at the close of the movement elicits significant ISC only in the latter occurrence. Given the importance of repetition in both musical structure (Margulis, 2012) and the formation of musical preferences over repeated listens (Szpunar et al., 2004), these findings highlight avenues for future research with a larger set of stimuli and variety of structural levels (e.g., song, section, phrase) over which musical content is repeated.

RC1 topographies for both stimuli are consistent with recently reported auditory RC1s (Cohen and Parra, 2016; Ki et al., 2016; Madsen et al., 2019; Kaneshiro et al., 2020) as well as other EEG components computed from responses to natural music (Schaefer et al., 2011; Sturm et al., 2015; Gang et al., 2017). While the phase-scrambled RC1 topography reported by Kaneshiro et al. (2020) was dissimilar to topographies of more musical stimuli, the present Control stimulus—in which intact music was phase scrambled and then scaled by its original amplitude envelope—elicits an RC1 topography that is broadly similar to that computed from intact music. In addition, while the Original stimulus elicited higher behavioral ratings and neural correlation than the Control stimulus, the Control also elicited significant ISC. The RC topography and ISC findings together distinguish the present Control stimulus from previous phase-scrambled controls without envelope scaling (Kaneshiro et al., 2020) and suggest that a music-like stimulus envelope, even one divorced from other attributes of music, drives correlated auditory responses.^7^ Future research can further characterize musical attributes driving neural correlation over a larger set of stimuli and stimulus manipulations.

The percentage of significant RC1 ISC windows for the Original stimulus (37.18%, Figure 3C) is broadly on par with previous reports related to film viewing (between approximately 21% and 33%, Dmochowski et al. (2012); 54.1%, Poulsen et al. (2017)). However, percentages of significant ISC for RC2 and RC3 were markedly lower during music listening compared to film viewing (Dmochowski et al., 2012). This finding is corroborated by sizeable drops in both the RCA coefficients (Figure 2B) and all-time ISC (Figure 3C). The drop in RCA coefficients has been reported previously by Kaneshiro et al. (2020) and motivated their decision to analyze only RC1 data. While we took an exploratory approach here in analyzing RC1–RC3 data (Dmochowski et al., 2012), our between-condition comparisons also ultimately focused on RC1. We recommend that standalone significance of single-RC ISC be carefully assessed before including those values in subsequent analyses, such as across-condition comparisons. EEG ISC is known to produce meaningful measures of brain states related to engagement.

Yet at its core, ISC is a group measure. Kaneshiro et al. (2020) have noted that the degree of abstraction embodied by music compared to movie narratives can impact the extent, and musical origin, of correlated responses: Compared to film viewing, music listeners may be more likely to focus on different stimulus attributes at different times, which would affect group measures such as ISC. Future work seeking individualized insights could assess one-against-all correlations of single participants, which have correlated with learning outcomes in a video-viewing study (Cohen et al., 2018). Stimulus-response correlation (SRC), the correlation of single EEG trials with time-varying stimulus attributes (Dmochowski et al., 2018), may be another way to index individual engagement during music listening. Forms of SRC have revealed significant group-level correlations across entire musical works (Gang et al., 2017; Kaneshiro et al., 2020). Future applications with time-resolved SRC—perhaps with participant-selected stimuli (Rickard, 2004; Grewe et al., 2007), which the paradigm enables—could prove useful in deriving single-listener time courses of engagement.

A time-varying neural measure of musical engagement suggests additional directions for future investigation. Our next work will augment present findings with the additional responses (ECG, respiratory activity, continuous behavioral reports) collected in the larger study. Other lines of future work can investigate how the extent of ‘narrative’ conveyed by music drives ISC. For instance, ‘program’ music conveying an extra-musical narrative, such as Prokofiev’s *Peter and the Wolf*, Op. 67 (Prokofiev, 1940), could bridge explicit narratives (Ki et al., 2016; Dmochowski et al., 2012), popular vocal music (Kaneshiro et al., 2020), and ‘absolute’ (i.e., non-representational) musical works used previously (Madsen et al., 2019) and here. Future investigations can also extend the use of time-varying ISC from our focus— on discrete time points corresponding to musical events identified a priori—to continuous measures that can be correlated in time with the ISC to elucidate connections between specific musical events and listener engagement. Such ‘second-order’ correlations could involve, for example, continuous listener reports of engagement (Broughton et al., 2019; Olsen et al., 2014), computationally extracted feature vectors (Lartillot and Toiviainen, 2007), or time-varying models of musical tension (Farbood, 2012).

Finally, EEG ISC can engender not only basic scientific insights, but also predictive insights for real-world application (Kaneshiro and Dmochowski, 2015). Taking a ‘hit song science’ (Pachet and Roy, 2008) approach, Leeuwis et al. (2021) showed that EEG alpha power synchrony could predict song popularity based on Spotify^8^ streams, when subjective ratings from the experimental sample could not. EEG ISC of small samples has also been used to predict population-level preferences across entire television commercials and in a time-resolved fashion over a television episode (Dmochowski et al., 2014). Current findings suggest that ISC may produce actionable within-*song* insights as well. Recent studies have proposed computational models for chorus and hook detection from audio (Van Balen et al., 2015) and used large-scale commercial data to model time-varying, within-song engagement using probabilities of Shazam^9^ queries (Kaneshiro et al., 2017) and Spotify skips (Montecchio et al., 2020). Thus, future investigations of temporally resolved ISC in conjunction with such reports have the potential to contribute to services aiding both music listeners and creators.

## Data Availability Statement

The data that support the findings of this study are openly available from the Stanford Digital Repository as part of the Naturalistic Music EEG Dataset— Elgar (NMED-E) at https://purl.stanford.edu/pp371jh5722.

## Abbreviations

(FDR): False Discovery Rate
(ISC): inter-subject correlation
(RCA): Reliable Components Analysis

## Acknowledgments

The authors thank Tysen Dauer and Daniel Abrams for helpful feedback on the manuscript, and Kunwoo Kim for contributions to exploratory data analysis. This research was funded by the Wallenberg Network Initiative: Culture, Brain Learning (BK, DTN, JB); the Patrick Suppes Gift Fund (BK, DTN); Army Research Laboratory W911-NF-10-2-0022 (JPD); and the Roberta Bowman Denning Fund for Humanities and Technology (BK, JB).

## Footnotes

1 The present analysis is not focused specifically on arousal. However, the data analyzed here are part of a larger study aimed to characterize both engagement and arousal during music listening (see § 2.4). Thus, the Elgar concerto was also chosen due in part to its self-selection by experimental participants in past research as an excerpt arousing strong emotions (Grewe et al., 2007) and use in a subsequent study of frisson, or ‘chills’ (Grewe et al., 2010).

2 https://www.audacityteam.org/

3 https://www.mathworks.com/

4 https://www.ee.columbia.edu/~dpwe/LabROSA/projects/

5 https://github.com/dmochow/rca

6 The RC1 topography for the Control stimulus was multiplied by −1 prior to plotting. This is due to a known arbitrary sign issue with RCA, which does not impact correlation.

7 We acknowledge that one phase-scrambled excerpt in Kaneshiro et al. (2020, Fig. 3C) did elicit significant ISC; however, we cannot directly compare findings given the differing component topographies across studies.

8 https://www.spotify.com/

9 https://www.shazam.com/

